# Relationship of *DUX4* and target gene expression in FSHD myocytes

**DOI:** 10.1101/2020.05.24.109710

**Authors:** Jonathan Chau, Xiangduo Kong, Nam Nguyen, Katherine Williams, Rabi Tawil, Tohru Kiyono, Ali Mortazavi, Kyoko Yokomori

## Abstract

Facioscapulohumeral dystrophy (FSHD) is linked to misexpression of the transcription factor, *DUX4*. Although DUX4 target gene expression is often readily detectable, analysis of *DUX4* expression has been limited due to its low expression in patient samples. Recently, single cell/nucleus RNA-sequencing was used to detect the native expression of *DUX4* for the first time, but important spatial relationships with its target gene expression was missing. Furthermore, dynamics of *DUX4* expression during myoblast differentiation has not been fully explored. In order to study the spatiotemporal relationship of *DUX4* and key target genes, we performed RNA FISH on immortalized FSHD2 patient skeletal muscle cells. Using two probe sets, *DUX4* transcripts were detected in 1-4% of myotubes after 3-day differentiation *in vitro*. We found that *DUX4* transcripts mainly localize as foci in one or two nuclei in a myotube compared to abundant accumulation of the target gene transcripts in the cytoplasm. Over a 13-day differentiation timecourse, *DUX4* expression without target gene expression significantly increased and peaked at day 7. Target gene expression correlates better with *DUX4* expression early in differentiation while the expression of target genes without detectable *DUX4* transcripts increases later. Consistently, shRNA depletion of DUX4-activated transcription factors, DUXA and LEUTX, specifically repressed a DUX4-target gene, *KDM4E*, later in differentiation, suggesting that following the initial activation by DUX4, target genes themselves contribute to the maintenance of downstream gene expression. Together, *in situ* detection of the *DUX4* and target gene transcripts provided new insight into dynamics of DUX4 transcriptional network in FSHD patient myocytes.

**Significance Statement:** FSHD is the third most common muscular dystrophy and is associated with upregulation of *DUX4*, a transcription factor, and its target genes. Although target genes are easily detectable in FSHD, low frequency *DUX4* upregulation in patient myocytes is difficult to detect, and examining the relationship and dynamics of *DUX4* and target gene expression without artificial overexpression of *DUX4* has been challenging. Using RNAScope with highly specific probes, we detect the endogenous *DUX4* and target gene transcripts *in situ* in patient skeletal myotubes during differentiation *in vitro*. Our study reveals a unique *DUX4* expression pattern and its relationship to the expression of target genes, and evidence for self-sustainability of the target gene network. The study provides important new insights into the FSHD disease mechanism.

## Introduction

Facioscapulohumeral dystrophy (FSHD) is an autosomal dominant muscular dystrophy initiating with progressive wasting of facial, shoulder, and upper arm musculature (1). It is one of the most common muscular dystrophies (1 in 8,333) (2). The majority of FSHD cases (>95%) are linked to monoallelic deletion of D4Z4 macrosatellite repeat sequences at the subtelomeric region of chromosome 4q (4qter D4Z4) (termed FSHD1 (MIM 158900)) (1, 3). Only one to ten D4Z4 repeats are found in the contracted allele in FSHD1 while 11∼150 copies are present in the intact allele. FSHD2 is the rare form of FSHD (<5% of cases) with no D4Z4 repeat contraction but exhibits the clinical phenotype identical to FSHD1 (4). Recent studies have found that the *SMCHD1* gene is mutated in >80% of FSHD2 cases (MIM 158901) (5) as well as in severe cases of FSHD1 (6, 7).

D4Z4 is a 3.3 kb repeat containing an open reading frame for the double-homeobox transcription factor *DUX4* gene (8-10). *DUX4* is essential during early embryogenesis but is subsequently silenced (11, 12). Only individuals with a 4qA haplotype with a polyadenylation signal sequence for the *DUX4* transcript distal to the last D4Z4 repeat express a full-length *DUX4* transcript (*DUX4fl*) and develop FSHD (13). Expression of *DUX4fl* is closely associated with FSHD, which strongly suggests that *DUX4* expression is critical for FSHD pathogenesis (10, 13, 14). Activation of many, if not all, DUX4 target genes has been observed in patient cells in multiple studies, supporting the significance of *DUX4fl* in FSHD. However, how dysregulation of any of these target genes directly contributes to the disease process is still under active investigation (9, 15-19).

Curiously, the *DUX4fl* transcript is expressed at extremely low levels and sometimes is not detectable (10, 17), and DUX4 protein is detectable only in <0.1% of patient muscle cells (10, 13, 14, 20). Furthermore, *DUX4fl* expression can occasionally be observed even in unaffected individuals (10, 17). Although overexpression of the recombinant DUX4 in in vitro myoblasts and in vivo in model organisms was shown to be toxic (21, 22), recent evidence indicates that the phenotype induced by the recombinant overexpression can differ from that of the endogenous DUX4 (23). Thus, there is a critical need to study the effect of the endogenous *DUX4* expression. However, assessment of the endogenous DUX4 and target gene expression in FSHD patient myocytes has been limited. Recently, we detected *DUX4* and target gene transcripts using single-nucleus RNA-seq (snRNA-seq) (24). Unlike the previous single cell RNA-seq of fusion-blocked myotubes (25), our isolation and analyses of nuclei from naturally fused multi-nucleated myotubes provided the first evidence that DUX4 target gene expression is much more wide-spread than *DUX4* transcription itself, which explains easier detection of the target gene transcripts rather than *DUX4* itself (9, 15-19). SnRNA-seq was highly instrumental in defining the different states of FSHD patient myocyte nuclei distinct from those of control myocyte nuclei. However, it failed to provide spatial relationship of individual nuclei and associated gene expression. In the current study, we examined the spatiotemporal relationship between the expression of *DUX4* and some of its major target genes in control and FSHD2 myocytes during differentiation using RNAScope, an *in situ* hybridization assay for RNA detection. We designed the probe set that maximizes the potential to detect *DUX4fl* and minimizes the crossreactivity with other isoforms and related genes. Our results provide the first direct visualization of *DUX4* and target gene activation in patient myocytes during differentiation as well as evidence for DUX4 target transcription factors controlling another DUX4 target gene. These results serve as an important basis for further understanding of the FSHD pathogenesis.

## Results and Discussion

### *DUX4* RNA accumulates in the nucleus of the FSHD myotubes

Multiple *DUX4* homologs and isoforms are known to be expressed in human myocytes (10). In particular, *DUX4s and DUX4c* were shown to be expressed more widely than *DUX4fl* even in control cells (Snider et al. 2010; Ansseau et al. 2009) (Supplemental Figure S1). Thus, to minimize crossreactivity and maximize the preferential detection of the full length *DUX4 (DUX4fl)* shown to be most relevant to FSHD, we custom designed an RNAScope probe set with the lowest possible number of ZZ probe pairs (“ZZ” represents a pair of RNAScope target probes) for the fluorescent detection system (Fig. 1A; 6ZZ). Interestingly, we observed the major *DUX4* transcript signal as foci in the nucleus using this set in primary FSHD myotubes (Fig. 1B) (24).

**Figure 1.**
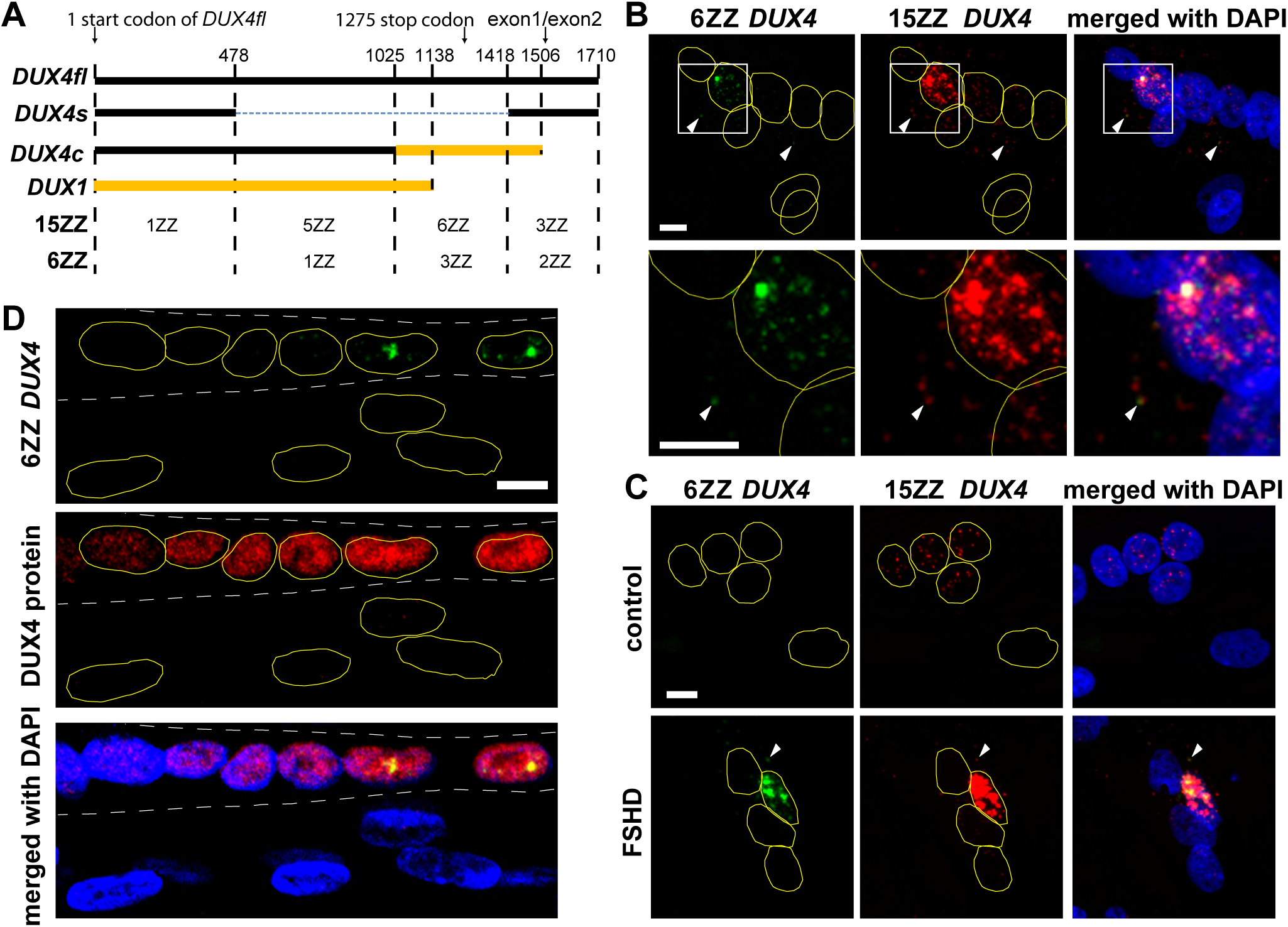
Specific detection of *DUX4* transcripts enriched in the FSHD myotube nucleus. **A.** Schematic diagram of mRNA transcripts for *DUX4fl*, the *DUX4s* isoform, *DUX4* homologs (*DUX4c* and *DUX1*), and the positions and numbers of individual ZZ pairs for the previously published RNAScope 15ZZ (26) and our 6ZZ probe sets. The black regions represent >99% homology to *DUX4fl*, which can be detected by corresponding ZZ pairs. The orange region in the 3’ half of *DUX4c* and DUX1 were confirmed not to crossreact with either 15ZZ or 6ZZ. As at least 3ZZ pairs are required for detectable signal, 15ZZ is capable of efficient detection of *DUX4s* and *DUX4c.* Our 6ZZ probe set was designed to preferentially detect *DUX4fl* and minimize the crossreactivity to *DUX4s* and *DUX4c*. Numbers indicate nucleotide number from the 5’ end of the transcripts. Nucleotide sequence comparison of four transcripts are shown in Supplemental Figure S1. **B.** RNAScope costaining with 15ZZ and 6ZZ probe sets in FSHD myotubes. Colocalization of the nuclear foci detected by both 15ZZ and 6ZZ. The lower panel is a magnification of the boxed region in the top panel. White arrowheads indicate the colocalization of both probe sets in the cytoplasm. Scale bar =10 µm. **C.** Additional nuclear foci in control and FSHD myotubes. Weak but distinct nuclear foci were observed in control cells with 15ZZ. No signal was observed with 6ZZ in control cells. Scale bar = 10 µm. **D.** Immunofluorescent staining of DUX4 protein (red) and RNAScope detection of *DUX4* transcript using 6ZZ probes (green) in primary FSHD myotubes at day 7 of differentiation. DAPI is in blue. White dashed lines indicate the boundary of a myotube with positive DUX4 antibody staining signal. Scale bar = 10 µm.

Unlike our probe set, previous studies using either a conventional FISH probe or another RNAScope probe set (ACDBio cat. no. 498541) detected *DUX4* transcript signals in the cytoplasm (26, 27). To address this apparent discrepancy, we performed costaining of the two RNAScope probe sets. The previous probe set was designed for the colorimetric DAB staining (26), which does not allow costaining. Thus, we remade the previous probe set compatible with fluorescent labeling and performed costaining with our new 6ZZ probe set. As expected, fluorescent labeling do not give as strong a signal as DAB (Fig. 1B) (26). Since the previous *DUX4* probe set contains 15 ZZ pairs (thus designated 15ZZ), the fluorescent signal is stronger than our 6ZZ probe set (Fig. 1B and C). Weaker staining by 6ZZ is due to fewer ZZ pairs in our probe set in order to minimize the potential crossreactivity to *DUX4s and DUX4c* (Fig. 1A). In the 15ZZ probe set, 3 and 5 ZZ pairs reside in two sub-regions of *DUX4fl* RNA shared by *DUX4s* and *DUX4c* transcripts, enabling 4ZZ and 6ZZ pairs to cross-hybridize *DUX4s* and *DUX4c*, respectively (Fig. 1A; 15ZZ). Consistent with this, 15ZZ shows some staining in control cells, in contrast to 6ZZ which has no significant signal (Fig. 1C). It is therefore possible that some of the signals detected by the 15ZZ probe set may come from that of *DUX4s* and/or *DUX4c*. Nevertheless, we found close colocalization of major fluorescent signals at nuclear foci by both probe sets, confirming the significant retention of *DUX4* transcripts in the nucleus (Fig. 1B). Some nuclear staining was also apparent in Figure 2C in the previous paper (26) though it was less clear due to the dark nuclear hematoxylin staining. We further confirmed the presence of DUX4 protein in the same myotube that contains the endogenous *DUX4* transcript-positive nuclei (Fig. 1D). Consistent with previous observations (19, 24, 28), expression of *DUX4* RNA transcripts in even one nucleus appears to be sufficient for DUX4 protein localization in most of the nuclei in the same myotube. Taken together, these results strongly support the specificity of the 6 ZZ probe set and indicate that the majority of *DUX4* transcripts are retained in the nucleus forming foci in FSHD myotubes.

**Figure 2.**
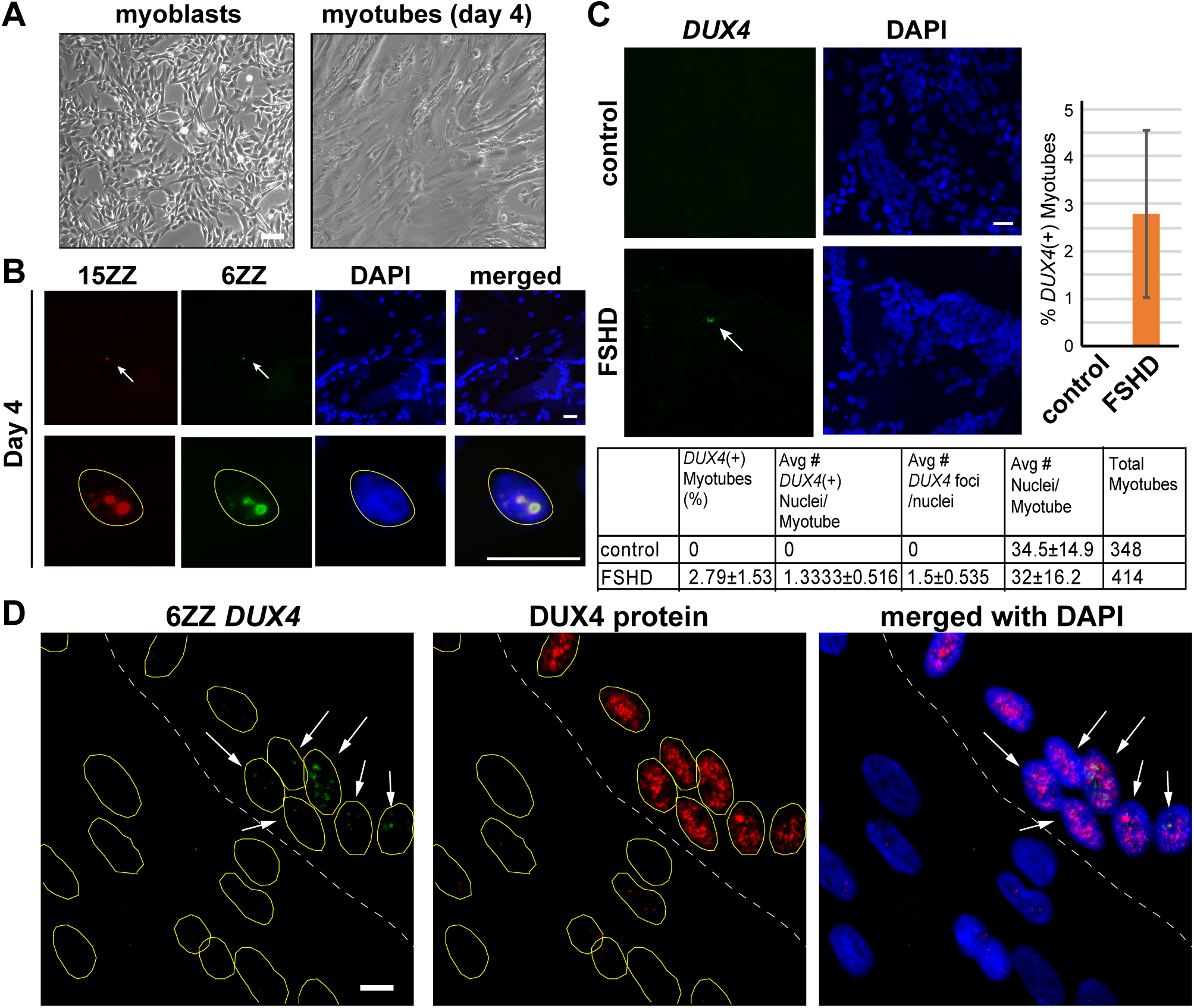
*DUX4* transcripts upregulated in FSHD myotubes. **A.** Bright field images of immortalized clonal FSHD2 myoblast differentiation on day 0 and day 4. Scale bar=100 µm **B.** RNAScope costaining of *DUX4* 15ZZ (red) and 6ZZ (green) probes in immortalized FSHD myotubes. White arrow indicates colocalization of probes. The lower panel is a magnification of the top panel. 15ZZ staining shows more foci and brighter signal similar to RNAScope staining in patient primary cells. Scale bar = 20 µm. **C.** Characterization of *DUX4* 6ZZ RNAScope in day 3 immortalized control and FSHD myotubes. *DUX4* staining usually appears in the nucleus and has around 1-2 bright foci. Around 2.8% of 414 FSHD2 myotubes contained a positive *DUX4* RNAScope signal. Scale bar = 20 µm. **D.** Detection of DUX4 protein by immunofluorescent staining and *DUX4* RNA by RNAScope. *DUX4* RNAScope (green) is combined with immunofluorescence using antibody against DUX4 (red) in immortalized FSHD myotubes at day 3 of differentiation. DAPI is in blue. White dashed line indicates the boundary of a myotube with positive DUX4 antibody staining signal. Arrow indicates nuclei with *DUX4* transcripts. Scale bar = 10 µm.

Cytoplasmic signals can also be observed by the 15ZZ probe set and weakly by ours in some FSHD myotubes with higher expression of endogenous *DUX4* (Fig. 1B, indicated by arrowheads). Since the cytoplasmic localization of the recombinant *DUX4* (*rDUX4*) transcript was observed previously (26), we also examined the localization of *rDUX4* transcript. Indeed, we found that the overexpressed r*DUX4* transcript is highly localized in the cytoplasm with much less nuclear localization (Supplemental Fig. S2). It is currently unclear whether this different RNA localization pattern is simply due to vast overexpression of the recombinant *DUX4* or possibly due to differences in template sequences and/or gene locations (ectopic vs. genomic). It is possible, for example, that untranslated regions, which are missing in the recombinant *DUX4* construct, may dictate the nuclear localization of the endogenous *DUX4* transcript. Nevertheless, the results revealed the significant difference between the endogenous and overexpressed recombinant *DUX4* transcript localization, highlighting the distinct retention in the nucleus of *DUX4* RNA transcribed from the endogenous locus.

### Quantification of *DUX4fl*-expressing nuclei in immortalized FHSD myotubes

To perform systematic analyses of *DUX4* and target gene transcripts, we established immortalized control and FSHD2 myoblast lines using *hTERT*, mutant *CDK4* and *CCND1* as described previously (29). We chose one particular patient myoblast sample for immortalization because of the significant expression of *DUX4* and target genes, which were closely characterized by snRNA-seq (24). After immortalization and surface marker isolation, several single clones were characterized and compared, and one of them that retained high proliferation and differentiation capabilities was chosen and used for the rest of the study. Using the immortalized FSHD myotubes on day 4 of differentiation (Fig. 2A), we observed similar nuclear foci of *DUX4* RNA by both 15ZZ and 6ZZ probe sets as in primary cells (Fig. 1), indicating that immortalization did not affect *DUX4* RNA expression and localization (Fig. 2B). We further quantified *DUX4* RNA signals detected by 6ZZ in control and FSHD myotubes. No signal was detected in control myotubes (N=348) (Fig. 2C). We observed *DUX4* RNAScope signals in one to two nuclei in ∼2.8% of FSHD2 myotubes on average (N=414) (Fig. 2C). Comparable *DUX4* RNAScope signal patterns and frequencies were observed in both primary parental and corresponding immortalized myotubes (Figs. 1 and 2) (24). Similar to the primary cells, we observed that all nuclei in a myotube are positive for DUX4 protein when some of the nuclei express *DUX4* RNA (Figs. 1D and 2D).

### Time course analyses of *DUX4* and target gene transcripts

To examine the relationship between DUX4 and its target transcripts, we designed probes for *LEUTX, KDM4E, ZSCAN4*, and *SLC34A2* transcripts known to be activated by DUX4 (19, 30, 31). We found specific expression of *LEUTX, KDM4E*, and *ZSCAN4* using our RNAScope probe sets in FSHD myotubes in a frequency similar to that of *DUX4* (Figs. 2C and 3A). We failed to detect any significant target gene signals in undifferentiated myotubes (data not shown). Unlike *DUX4*, however, these target gene transcripts, if expressed, accumulate abundantly in the cytoplasm (Fig. 3A). The probe set for *SLC34A2* detected strong cytoplasmic signals in FSHD myotubes (Fig. 3A). However, some non-specific staining was seen in control cells, and thus the probe set appears to be not completely specific to FSHD-induced *SLC34A2* RNA (data not shown). Thus, we eliminated this probe from further analyses. We confirmed the presence of LEUTX protein in all the nuclei in the same myotube with *LEUTX* RNA signal, confirming the correlation of RNA and protein expression (Fig. 3B, middle). We observed one myotube with weak LEUTX protein staining and no RNA (Fig. 3B, bottom). It is unclear whether this represents a myotube with residual LEUTX protein after mRNA transcription was ceased. With these results, we chose *LEUTX* and *KDM4E* transcripts for further time course analyses.

**Figure 3.**
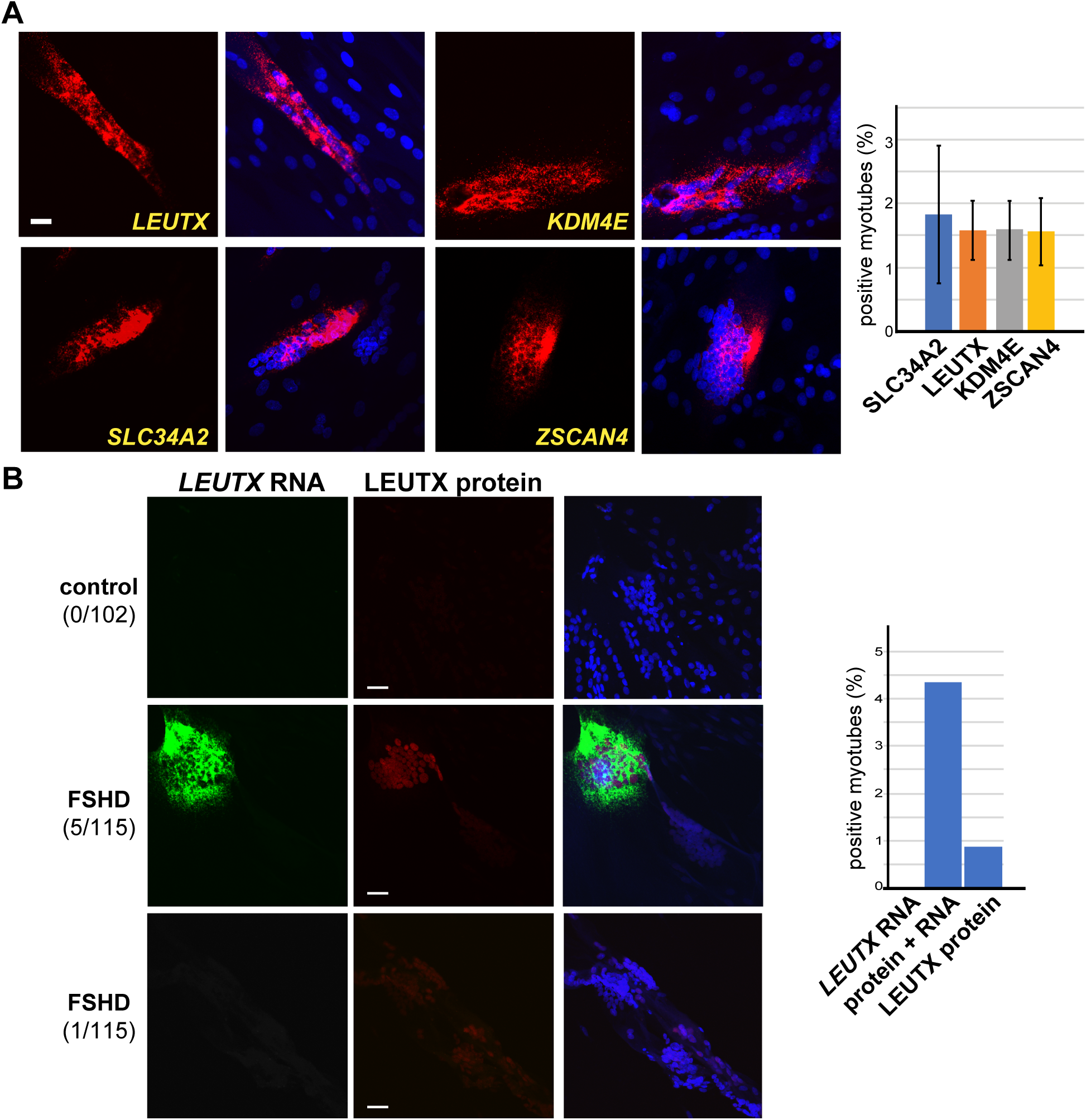
Comparison with DUX4 target gene expression. **A.** RNAScope staining of DUX4 target genes *LEUTX, KDM4E, SLC34A2*, and *ZSCAN4*. Unlike *DUX4*, staining of target genes is spread along the myotube and in the cytoplasm. Approximately 2% of myotubes showed positive RNAScope target gene signal. Scale bar = 20 µm. **B.** Costaining of *LEUTX* RNAScope (green) and LEUTX protein immunofluorescence staining (red) in control and FSHD myotubes on differentiation day 6. The middle panel shows a typical example of RNA and protein colocalization while the bottom panel shows an example of weak protein staining without RNA. Numbers of myotubes with corresponding patterns in the total myotubes counted are shown in parentheses. Percentages of myotubes with *LEUTX* RNA only, *LEUTX* RNA and protein, or LEUTX protein only are shown. Scale bar = 20 µm.

While the upregulation of DUX4 target genes in FSHD is relatively well established, how *DUX4* activation results in target gene expression during human myoblast differentiation has not been studied at the single cell level in the context of the natively fused myotubes. Previous work to understand this question includes single cell RNA-seq done on fusion-inhibited myocytes (25) and our recent single-nucleus RNA-seq (24). However, neither study was able to address dynamics of *DUX4* and target gene expression in the intact myotube. Thus, we performed triple staining for endogenous *DUX4, LEUTX, and KDM4E* expression to investigate their localization *in situ* during differentiation from day 3 through day 13 (Fig. 4 and Supplemental Figure S3). Because the number of all *DUX4* and target gene-expressing myotubes is limited (typically ∼2% for days 3-5 and ∼5% for later days of the entire myotube population on a coverslip), obtaining statistically significant quantification data is challenging. Nevertheless, with multiple experimental replicates, we were able to observe a reproducible trend for *DUX4* and target gene expression kinetics during differentiation (Fig. 4B). Previously, we followed the increase of DUX4 target gene induction up to 5 days of differentiation by RNA-seq (24). In the current study, we observed that the number of myotubes expressing target genes (*LEUTX* and/or *KDM4E* with or without *DUX4)* has a tendency to increase throughout the duration of a 13-day time course (Fig. 4C, pink and blue). Increasing trajectory of *DUX4* and target gene-positive myotubes argue against the previous suggestion that *DUX4* expression leads to immediate cell death (19, 32). We also found that the number of myotubes with *DUX4* only expression (without any target gene expression) increases significantly starting at day 3 and peaks at day 7 (Fig. 4B and C green), indicating that there are two states of *DUX4*-positive myotubes (with or without downstream gene activation). The results raise the possibility that an additional factor(s) may be involved in efficient downstream target gene activation.

**Figure 4.**
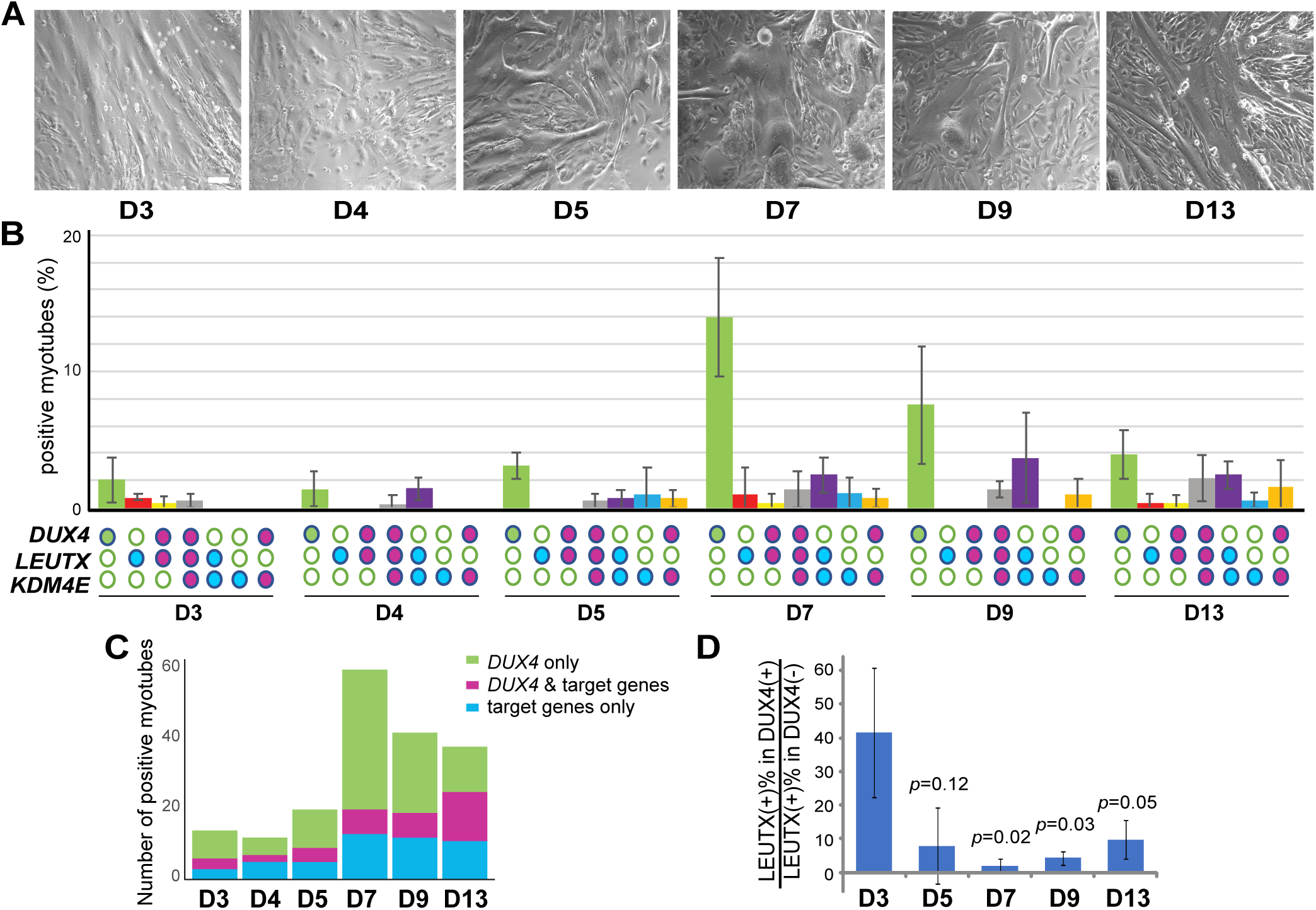
Dynamic relationship between *DUX4* and target gene expression during myotube differentiation. **A.** Bright field images of immortalized FSHD myoblast differentiation into myotubes on days 3-13. Around day 5, we find greater myotube detachment. Scale bar =100 µm **B.** Quantification of time course RNAScope triple staining of *DUX4, LEUTX*, and *KDM4E* transcripts. Error bars indicate the standard deviation from the mean of three independent experiments. The actual number counts are shown in Supplemental Figure S3. **C.** Frequencies of myotubes expressing *DUX4 only, DUX4* and target genes, or target genes only change during differentiation. Replotting the data in (B) for the number of myotubes containing *DUX4* transcripts only, transcripts of DUX4 plus target gene(s), or transcripts of target gene(s) only as indicated. Y-axis is the total number of positive myotubes counted. **D.** Frequency of *LEUTX* expression in *DUX4*-expressing myotubes decreases later in differentiation. The ratios between the percentage of *LEUTX*-expressing (LEUTX(+)) myotubes in the entire *DUX4*-expressing (DUX4(+)) myotubes and the percentage of LEUTX(+) myotubes in myotubes with no *DUX4* expression (*DUX4*(-)) at different days after differentiation were calculated based on the RNAScope data in (B). At day3, the frequency of DUX4(+) myotubes to co-express *LEUTX* is ∼20-60 fold higher than the frequency of *DUX4*(-) myotubes to express LEUTX. Later in differentiation (days 7 and 9), the ratios drop significantly, indicating that the frequency of *LEUTX* expression without *DUX4* expression becomes more significant. The error bars represent the standard deviation of the mean of 3 independent experiments. The significance was evaluated by student t-test. The *p* values were calculated comparing to day 3. *p* < 0.05 was considered significant.

Unexpectedly, the number of myotubes expressing two target genes (*LEUTX* and/or *KDM4E*) without *DUX4* increased later in differentiation (Fig. 4C blue), suggesting their continued upregulation with no *DUX4* transcripts present. Consistent with this, the frequency of *LEUTX* coexpression with *DUX4* decreases significantly later in differentiation (Fig. 4D). These results strongly suggest that for those myotubes in which the target genes are activated (initially by DUX4), their expression may continue even when *DUX4* expression is not maintained. Taken together, the results reveal some discordance between *DUX4* and target gene expression, suggesting the possible presence of additional regulatory mechanisms for the DUX4 gene network activation in FSHD patient myocytes.

### *KDM4E* expression is regulated by DUXA and LEUTX

The above results reveal coexpression of DUX4 target genes in the same myotubes with no detectable *DUX4* (Fig. 4B and C). In particular, the number of *KDM4E* expressing myotubes increases even after *DUX4* expression peaks at day 7 (Fig. 5A). Coexpression of *KDM4E* and *LEUTX* without *DUX4* increases later in differentiation (Fig. 4C, blue) and *KDM4E* transcript expression correlates better with expression of *LEUTX* than *DUX4* (Fig. 5B). Costaining of LEUTX protein with *KDM4E* transcripts on day 6 revealed that all observed *KDM4E* staining co-localized with LEUTX protein staining in the same myotubes (Supplemental Fig. S4). Motif analyses revealed that the promoter region of the *KDM4E* gene contains not only the DUX4 binding motif, but also multiple sites of the putative LEUTX binding motif (Supplemental Fig. S5) (33, 34). These results raise the possibility that LEUTX may be involved in *KDM4E* upregulation. We recently found that DUXA, another DUX4 target, plays a significant role in upregulating *LEUTX* later in differentiation (24). Thus, we depleted LEUTX and DUXA by shRNAs (Fig. 5C). Because of the difficulty detecting DUXA and LEUTX proteins by western blot, depletion efficiency was confirmed by RT-qPCR (Fig. 5D). Although depletion efficiency is at a comparable level on day 4 and day 6 of differentiation, LEUTX and DUXA depletion specifically repressed *KDM4E* expression on day 6, but not day 4. While we do not detect DUXA motifs in the promoter of *KDM4E*, we cannot preclude DUXA binding given the similarity of its motif to that of DUX4. Taken together, these results indicate that LEUTX (and DUXA either directly or indirectly through LEUTX upregulation) promotes *KDM4E* expression following the initial activation by DUX4 (Fig. 5E). The results support our hypothesis that once activated, the DUX4 target genes may in part self-sustain their expression (24).

**Fig. 5.**
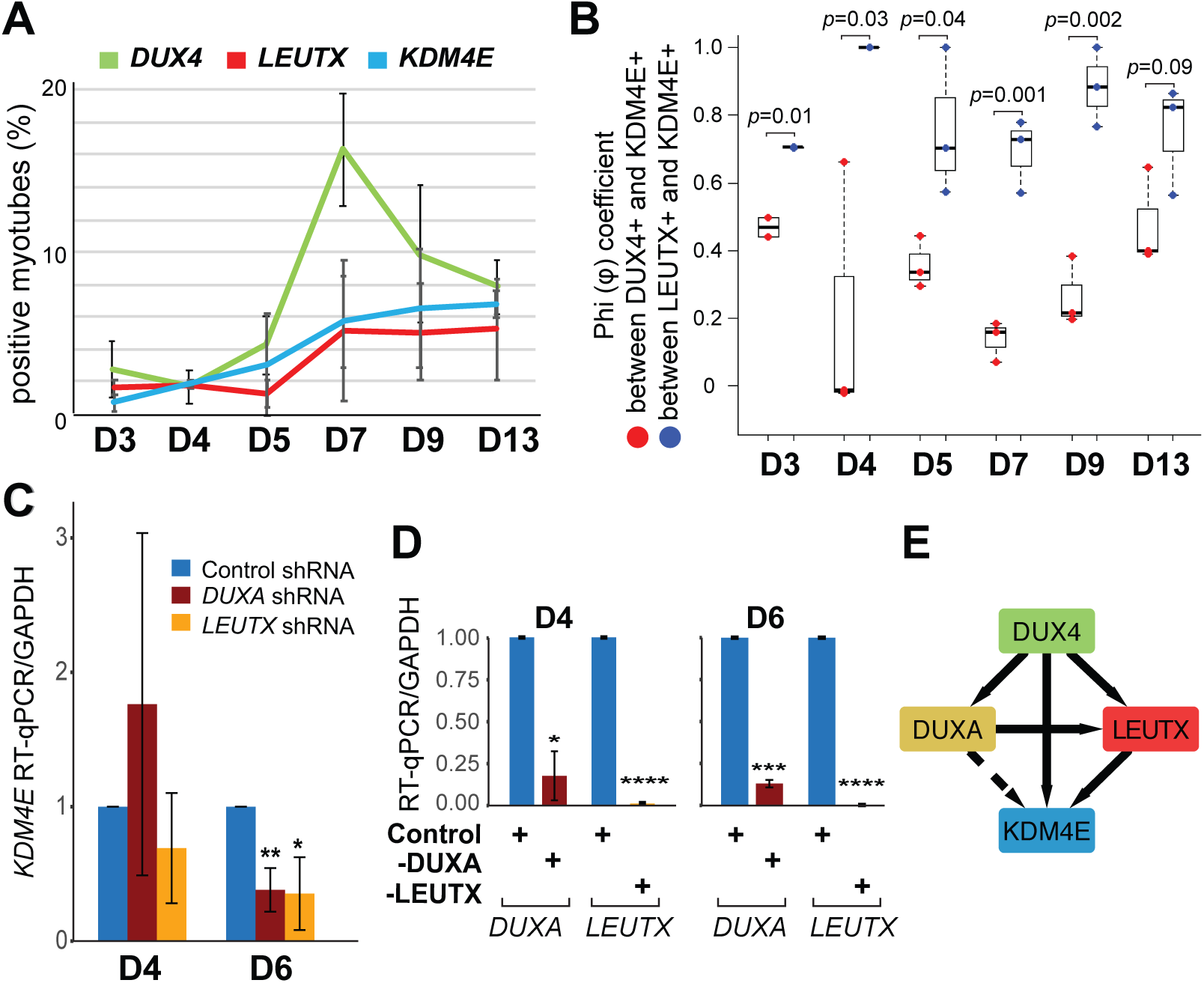
*KDM4E* expression is regulated by DUX4 target transcription factors. **A.** Individual timecourse RNAScope analysis of *DUX4, LEUTX*, and *KDM4E* in FSHD myotubes for the indicated days of differentiation from Figure 4B. These include all the positive myotubes for each transcript. **B.** Comparison of correlation between *KDM4E* and *DUX4/LEUTX* expression based on the RNAScope data in Figure 4B. The high φ values indicate strong positive relationship between *KDM4E* and *LEUTX* RNA expression for the entire duration of the timecourse, with a perfect positive correlation (φ=1) on day 4. After that, the φ values decrease but are more significant than those between *KDM4E* and *DUX4* expression until day 13. P-values (unpaired, two-tailed t test) are for the correlation differences of indicated genes at the same day. **C.** The effect of *DUXA* or *LEUTX* depletion on *KDM4E* expression in FSHD myotubes.. *KDM4E* gene expression in control, *DUXA*, or *LEUTX* shRNA-treated myotubes on day 4 and day 6 was measured by RT-qPCR. The signal was normalized to *GAPDH*. The bar graph shows the average fold change of three independent experiments for each knockdown compared to the control shRNA-treated cells on day 4 or day 6 as indicated. Student t-test was used to calculate p-values. * p-value <0.05, ** p-value <0.01. **D.** Analysis of depletion efficiency of *DUXA* and *LEUTX* shRNA by RT-qPCR. Student t-test was used to calculate p-values for expression of *DUXA* or *LEUTX* in corresponding shRNA-treated myotubes compared to shRNA control on days 4 and 6 of differentiation. * p-value<0.05, *** p-value<0.001, **** p-value<0.0001. **E.** A schematic diagram depicting the proposed relationship between DUX4 and the target genes. DUX4 expression is important for induction of *DUXA, LEUTX* and *KDM4E* during early FSHD myoblast differentiation. Once expressed, LEUTX contributes to *KDM4E* upregulation. DUXA activates *LEUTX* and *KDM4E* (directly and/or possibly through LEUTX).

Detection of the endogenous *DUX4* expression and its relationship with its target gene expression in patient myocytes has been technically challenging due to the low frequency of *DUX4* expressing cells. Here we use a custom-designed RNAScope probe set to maximize the detection of the pathogenic full-length *DUX4* transcript and to analyze its localization and relationship with its downstream target genes in differentiating FSHD myotubes. The use of RNAScope ((26) and the current study) provides a complementary tool to single-cell/nucleus RNA-sequencing, to understand FSHD pathogenesis with high spatiotemporal resolution. Our results provide snapshots of FSHD-induced gene expression in patient myocytes during differentiation, revealing that (1) the endogenous *DUX4* RNA mainly accumulates in the nucleus; (2) *DUX4* expression increases over a week of differentiation, suggesting that it is not immediately toxic; (3) *DUX4* and target gene expression are not always concordant, identifying different states of *DUX4*-positive myotubes; and (4) once activated by DUX4, target genes by themselves can contribute to sustaining DUX4-induced gene expression changes, and that LEUTX, a primate-specific transcription factor like DUX4, is involved in this process. Taken together, our findings provide new insight into gene expression changes in patient myocytes, and evidence for an additional layer of regulation to the DUX4-induced transcriptional network.

## Methods

### Cell culture and differentiation

Primary and immortalized control and FSHD2 skeletal myoblast cells were grown in high glucose DMEM (Gibco) supplemented with 20% FBS (Omega Scientific, Inc.), 1% Pen-Strep (Gibco), and 2% Ultrasor G (Crescent Chemical Co.). Primary control and FSHD2 myoblasts (24) were immortalized using hTERT with p16INK4a-resistant R24C mutant CDK4 (mtCDK4) and Cyclin D1 as previously described (35). After immortalization, CD56-positive cells were selected by magnetic-activated cell sorting conjugated with anti-CD56 antibody (130-050-401, MiltenyiBiotec). Single cell clones were isolated by FACS sorting into 96 well plates. Control and FSHD2 clones were chosen for the experiments based on normal doubling time and high differentiation index. Myoblast differentiation was induced as previously described (29). Briefly, cells were plated at a seeding density of ∼2.5 × 10^5^ cells/ml in 0.5 ml of growth medium in each well of a 24-well dish. Approximately 12-16 hr later differentiation was induced using high glucose DMEM medium supplemented with 2% FBS and ITS supplement (insulin 0.1%, 0.000067% sodium selenite, 0.055% transferrin, 51300044 Invitrogen). Fresh differentiation medium was changed every day.

### Antibodies and cDNA clone

Immunofluorescence was performed using rabbit polyclonal antibodies specific for DUX4 (ab124699, Abcam) and LEUTX (PA5-59595, Thermofisher). The recombinant DUX4 expression plasmid (pCS2-mkgDUX4) was a gift from Dr. Stephen Tapscott (Addgene plasmid # 21156) (36).

### RNAScope probe design

The following RNAScope probes (Advanced Cell Diagnostics, Inc.) were used: *LEUTX* (Hs-LEUTX-C2, Cat No. 547251-C2), *KDM4E* (Hs-KDM4E-C3, Cat No. 556121-C3), *SLC34A2* (Hs-SLC34A2-C3, Cat No. 407101-C3), *ZSCAN4* (Hs-ZSCAN4-C2, Cat No. 421091-C2) and 15ZZ *DUX4* (Hs-DUX4-No-XMm-C3, Cat No. 498541-C3). The 6ZZ *DUX4fl* probe set (HS-DUX4-O6-C1, Cat No. 546151) was specifically designed for *DUX4fl* (NM_001306068.2), with only 1 or 2 ZZ pairs residing in the regions shared with *DUX4s* or *DUX4c* (at least 3ZZs are needed for signal detection) (Fig. 1A).

### RNAScope hybridization

Cells were grown and differentiated on cover slips in 24-well plates. Cells were washed twice with PBS, then fixed with 10% neutral buffered formalin (NBF) for 30 min at room temperature (RT), and finally dehydrated with 50%, 70%, and 100% ethyl alcohol gradients for 1 min each at room temperature. Cells were then rehydrated with 70% and 50% ethyl alcohol gradients for 1 min each and finally treated with PBS for 10 min. Cells were then treated with hydrogen peroxide (Cat No. 322335 ACDBio) and protease III (Cat No. 322337 ACDBio) at RT for 10 min each and washed with PBS. The probe sets for *DUX4, LEUTX, KDM4E* were then added in a 50:1:1 ratio for 2 h at 40°C within a humidity control chamber. Probe sets are optimized for different fluorescent channels (C1, C2, and C3, respectively). RNAScope multiplex signal amplification (Cat No. 323110 ACDBio) were applied sequentially and incubated in AMP 1, AMP 2, AMP 3 for 30, 30, 15 min respectively at 40°C within the humidity control chamber. Before adding each AMP reagent, cells were washed twice with RNAScope washing buffer (Cat No. 310091 ACDBio). Signal was developed with the Fluorescent Detection Reagents (Cat No. 323110 ACDBio) and TSA Plus Fluorophores diluted 1:1500 in RNAScope TSA Buffer (Cat No. 322810 ACDBio). To develop C1 signal, samples were incubated in HRP-C1, TSA Plus Fluorescein (cat. No. NEL741001KT PerkinElmer), HRP Blocker for 15, 30, 15 min, respectively, at 40°C within the humidity control chamber. Before adding each reagent, cells were washed twice with RNAScope washing buffer. For C2 and C3 costaining, TSA Plus Cy 3 and Cy 5 were used, respectively (cat no. NEL744001KT and NEL745001KT PerkinElmer). Cells were then counterstained with DAPI (Cat No. 320858 ACDBio) for 30s at RT and washed twice with PBS. Samples were then dried and mounted onto microscope slides with ProLong Diamond Antifade Mountant (Cat No. P36961 ThermoFisher).

### Immunofluorescent co-staining with RNAScope

Cells were grown and differentiated on coverslips in 24-well plates. For costaining of DUX4 antibody and DUX4 RNAScope probes, cells were fixed with 10% NBF for 30 min at RT and then extracted with 0.5% Triton X-100 in PBS. Primary and secondary antibodies were diluted in SNBP (1XPBS /0.02% saponin /0.05% NaN_3_ /1% BSA), containing 1% horse serum and 0.05% gelatin. Samples were incubated in primary antibody for 30 min at 37°C followed by three PBST (PBS with 0.05% Tween20) washes. Coverslips were incubated in secondary antibody for 30 min at 37°C followed by three PBST washes. Cells were then dehydrated with 50%, 70%, and 100% ethyl alcohol gradients for 1 min each at RT and continued to be processed for RNAScope hybridization as described above. Then coverslips were counterstained with DAPI, washed with dH_2_O and mounted with Prolong Diamond Antifade Mountant. For costaining of LEUTX protein and *LEUTX* RNA, the RNAscope protocol was done before the immunofluorescence. Before DAPI counterstaining in the RNAscope procedure, coverslips were blocked in PBST, containing 2% BSA and 10% Milk for 45 min. Primary and Secondary antibodies were diluted in the blocking buffer. Samples were incubated in primary antibody for 2h at RT followed by three PBST washes. Coverslips were then incubated in secondary antibody for 1h at RT followed by three PBST washes. Then, coverslips were counterstained with DAPI, washed with PBST and mounted with Prolong Diamond Antifade Mountant.

### Fluorescent image acquisition, quantification and statistical analysis

Images were acquired with a Zeiss LSM510 confocal laser microscope. On the 12 mm diameter coverslip, 5 horizontal sections were observed and myotubes were counted. Three replicates were done of each staining experiment and error bars were calculated using standard deviation from the mean.

### ShRNA depletion and RT-qPCR

Lentiviruses carrying shRNA plasmids (MISSION shRNA, Sigma-Aldrich) for each DUX4 target gene: *DUXA* (5’-CTAGATTACTTCTCCAGAGAA-3’, TRCN0000017664), *LEUTX* (5’-CCTGGAATCTCTGATGCAAAT-3’, TRCN0000336862), and an shRNA control (SHC002) were made in 293T cells using Lipofectamine P3000. The cells were transfected with 2 µg of shRNA plasmids, 1.5 µg of pCMV plasmids, and 0.5 µg of pMP2G plasmids (37). The media was changed after 24 hours. The lentiviruses were harvested at 48 hour and 72 hour post-transfection. FSHD2 immortalized myoblasts were infected twice at 32 hour and 8 hour prior to differentiation. The myoblasts were selected with puromycin. The RNA was extracted using RNeasy kit (Qiagen, Cat No. 74134) at days 4 and 6. Around 16 ng of RNA was converted to cDNA using SuperScript IV VILO Master (Thermofisher, Cat No. 11756050) and then used for RT-qPCR analysis. PCR primers are listed in Supplemental Table S1.

### Statistical Analyses

Microsoft Excel software was used to perform statistical analyses on data from three independent experiments. The statistical significance is determined by the Student’s t-test. P < 0.05 is considered of statistical significance. The error bars denote standard deviation. Phi (φ) coefficient (analytically equivalent to Pearson’s correlation for binary data) is calculated following the previous paper (38). φ takes on values ranging between +1 and -1. The following points are the accepted guidelines for interpreting the correlation coefficient: “φ=1”, “1>φ>0.5”, “0.5>φ>0.3” and “0.3>φ>0.1” indicate perfect, strong, moderate and weak positive relationship respectively; “0.1>φ>-0.1” indicates no relationship; while “φ=-1”, “-1<φ<-0.5”, “-0.5<φ<-0.3” and “-0.3<φ<-0.1” indicate perfect, strong, moderate and weak negative relationship respectively.

## Acknowledgement

The authors wish to acknowledge the support of the Chao Family Comprehensive Cancer Center Optical Biology Core (LAMMP/OBC) Shared Resource. We would also like to thank Dr. Naohiro Hashimoto (National Center for Geriatrics and Gerontology, Japan) for technical advice and Kevin Cabrera for helpful comments on the manuscript. This work was supported in part by National Institutes of Health (P01NS069539 to R.T., and R01AR071287 to K. Y. and A. M.).

## Figure legends

**Supplemental Table S1.**
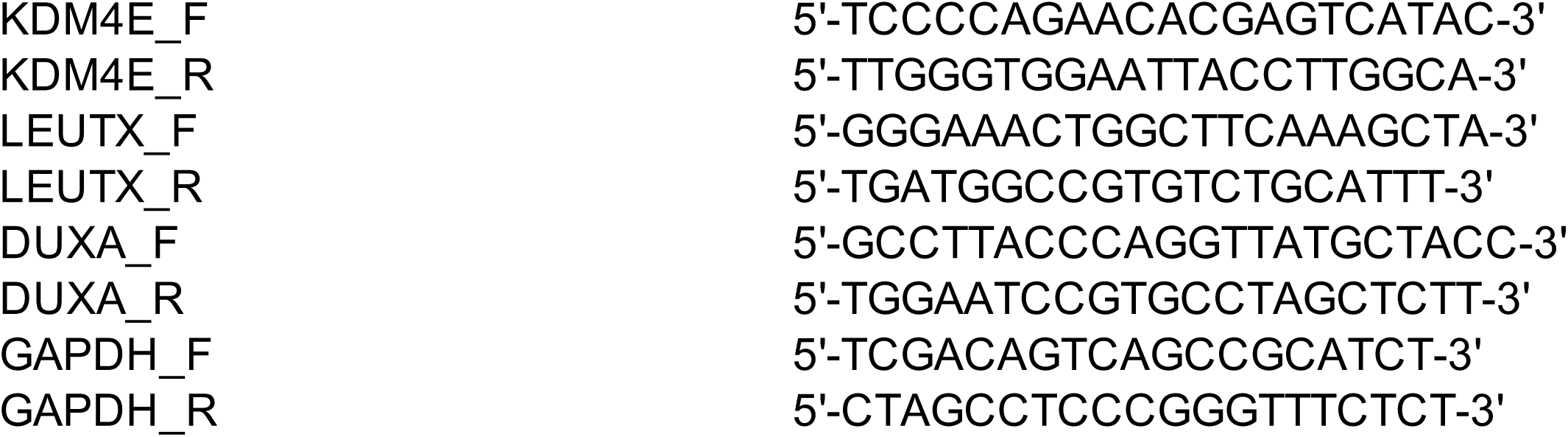
PCR primers.

**Supplemental Figure S1.**
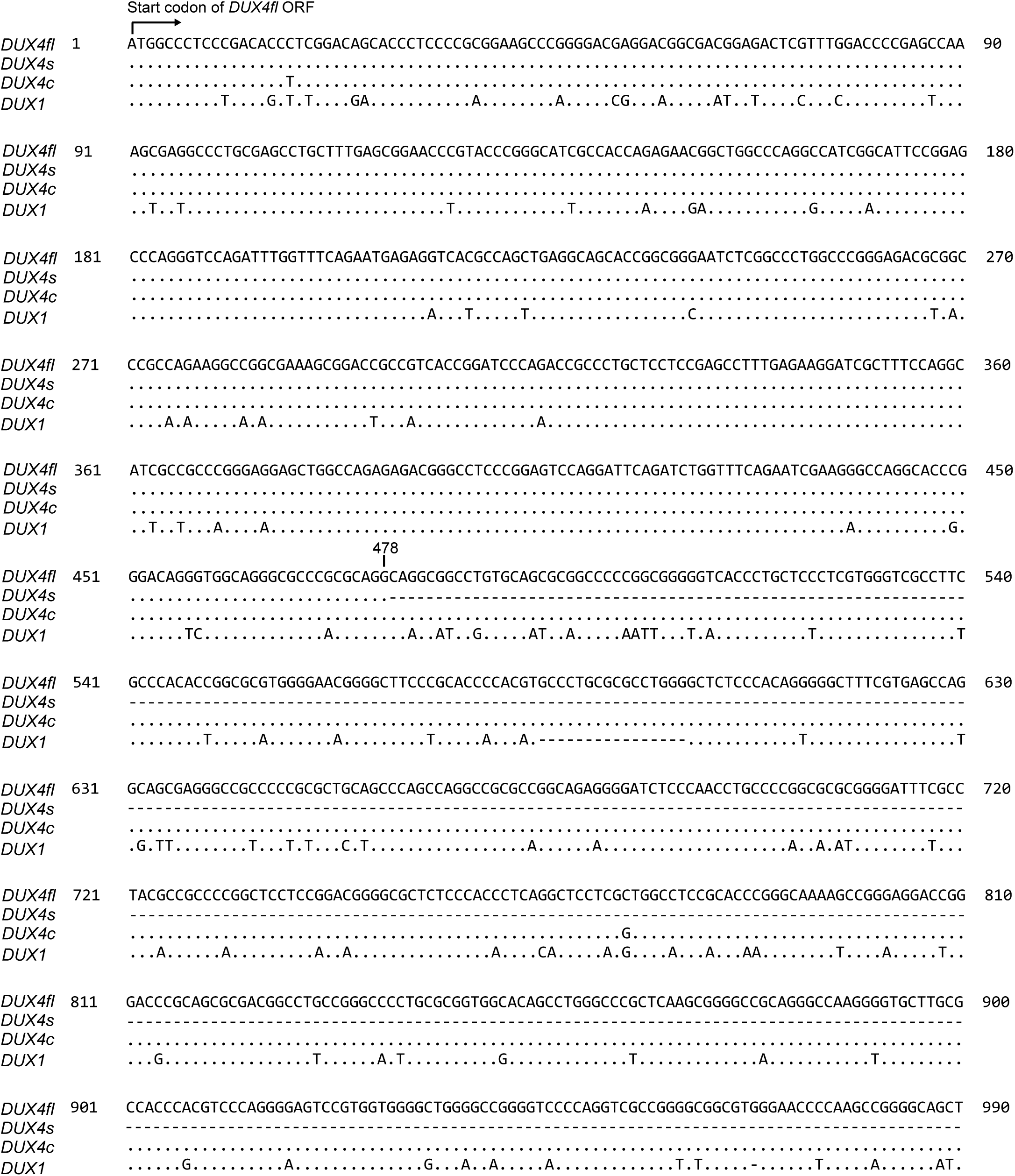
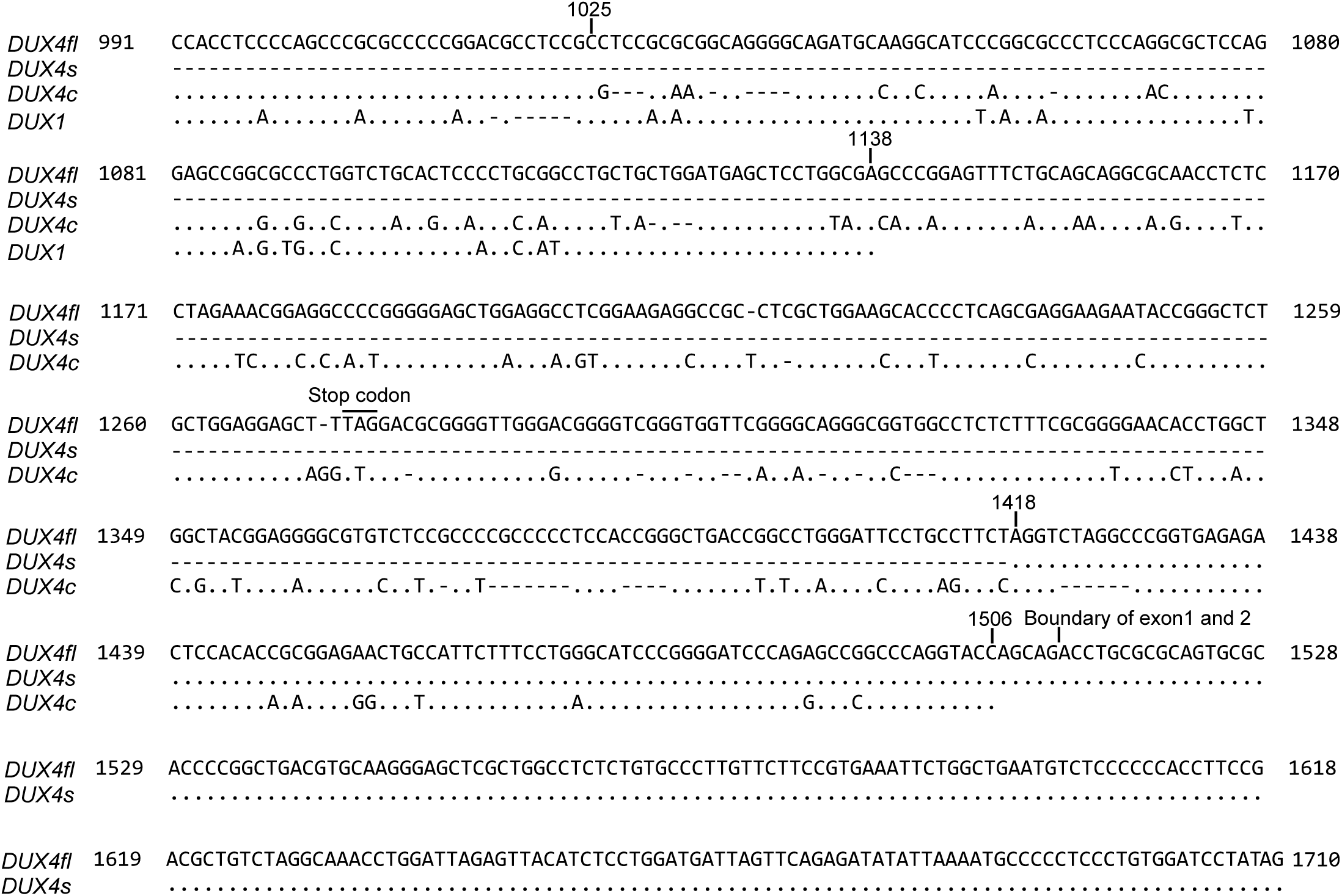
Sequence comparison of *DUX4fl, DUX4s, DUX4c* and *DUX1* transcripts. *DUX4s, DUX4c* and *DUX1* transcripts alignments are shown in relation to *DUX4fl*. Identities are displayed as dots (.), with mismatches displayed as single letter abbreviations. Dash marks (-) indicate deleted regions. The start codon, stop codon and exon 1/exon 2 boundary of *DUX4fl*, as well as the regions boundaries in Figure 1A were labeled on top of *DUX4fl* sequence. Nucleotide numbering was based on *DUX4fl* transcript with accession number NM_001306068.2 in the NCBI database.

**Supplemental Figure S2.**
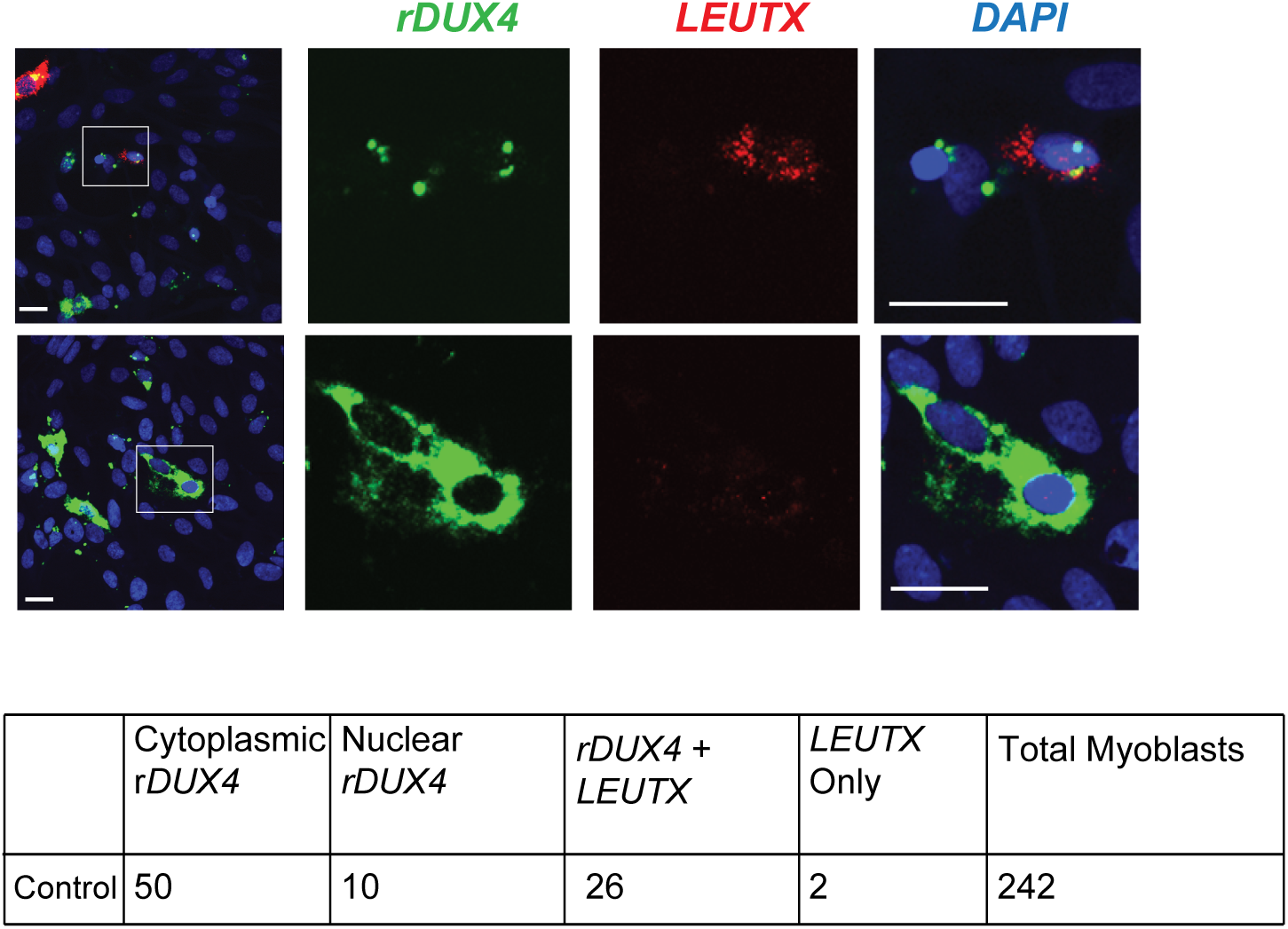
RNAScope analysis of the transiently transfected recombinant *DUX4* (*rDUX4*) and the endogenous *LEUTX* RNA expression in immortalized control myoblasts. Example images of different *rDUX4* RNA localization patterns are shown. The boxes in the first column indicate where the image is zoomed in. Table underneath shows the quantification of the number of myoblasts with different patterns of *rDUX4* and *LEUTX* RNA signals. Majority of *rDUX4*-transfected myoblasts exhibited strong cytoplasmic *rDUX4* RNA signals. *LEUTX* expression is not always seen in *rDUX4*-expressing cells. Scale bar = 20 μm.

**Supplemental Figure S3.**
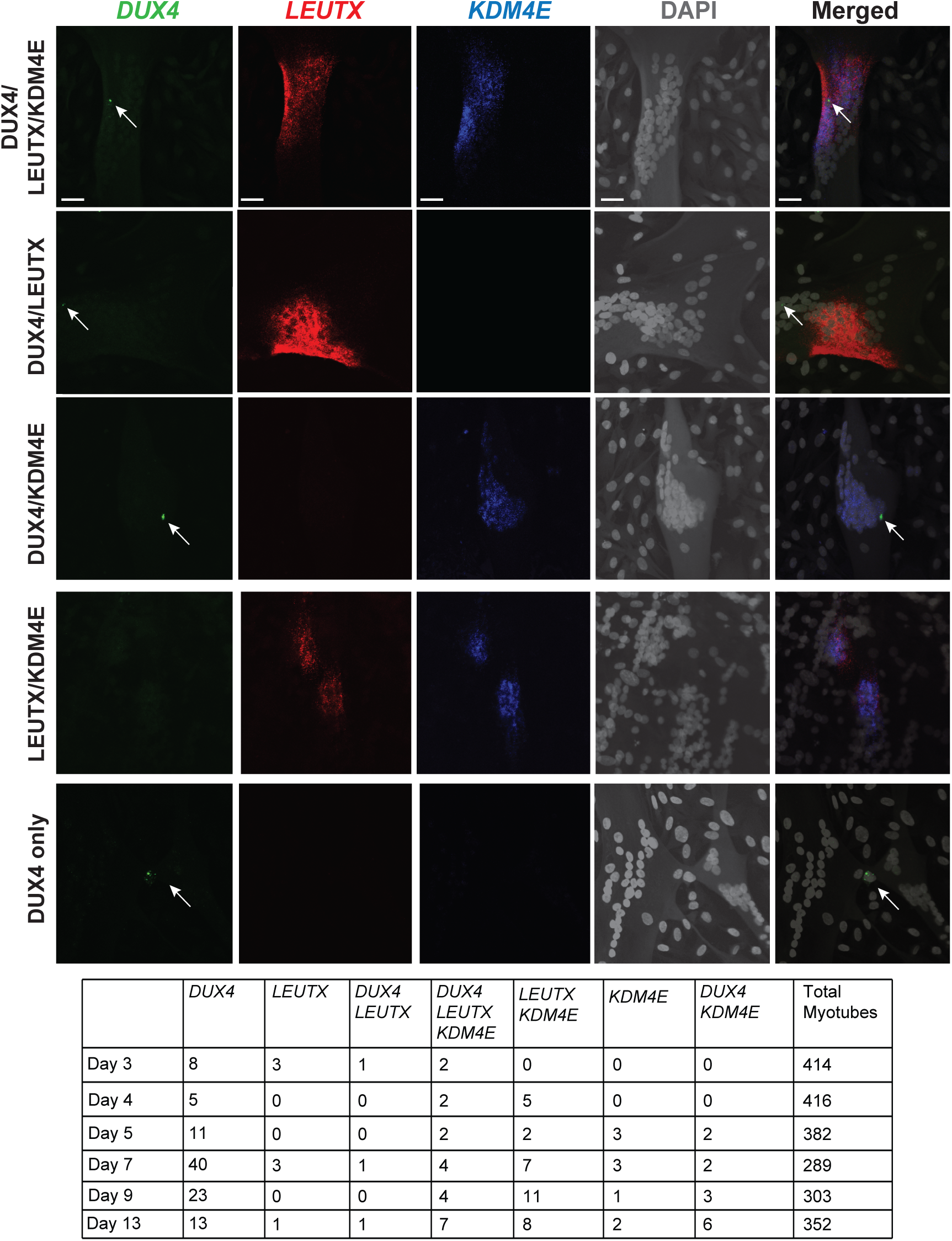
Representative images of myotubes triple stained with RNAScope for *DUX4, LEUTX*, and *KDM4E* RNA during differentiation providing data for Figure 4. White arrows indicate *DUX4* transcript-positive nuclei. The table shows the total counts from the three replicates. Scale bar = 20 μm.

**Supplemental Figure S4.**
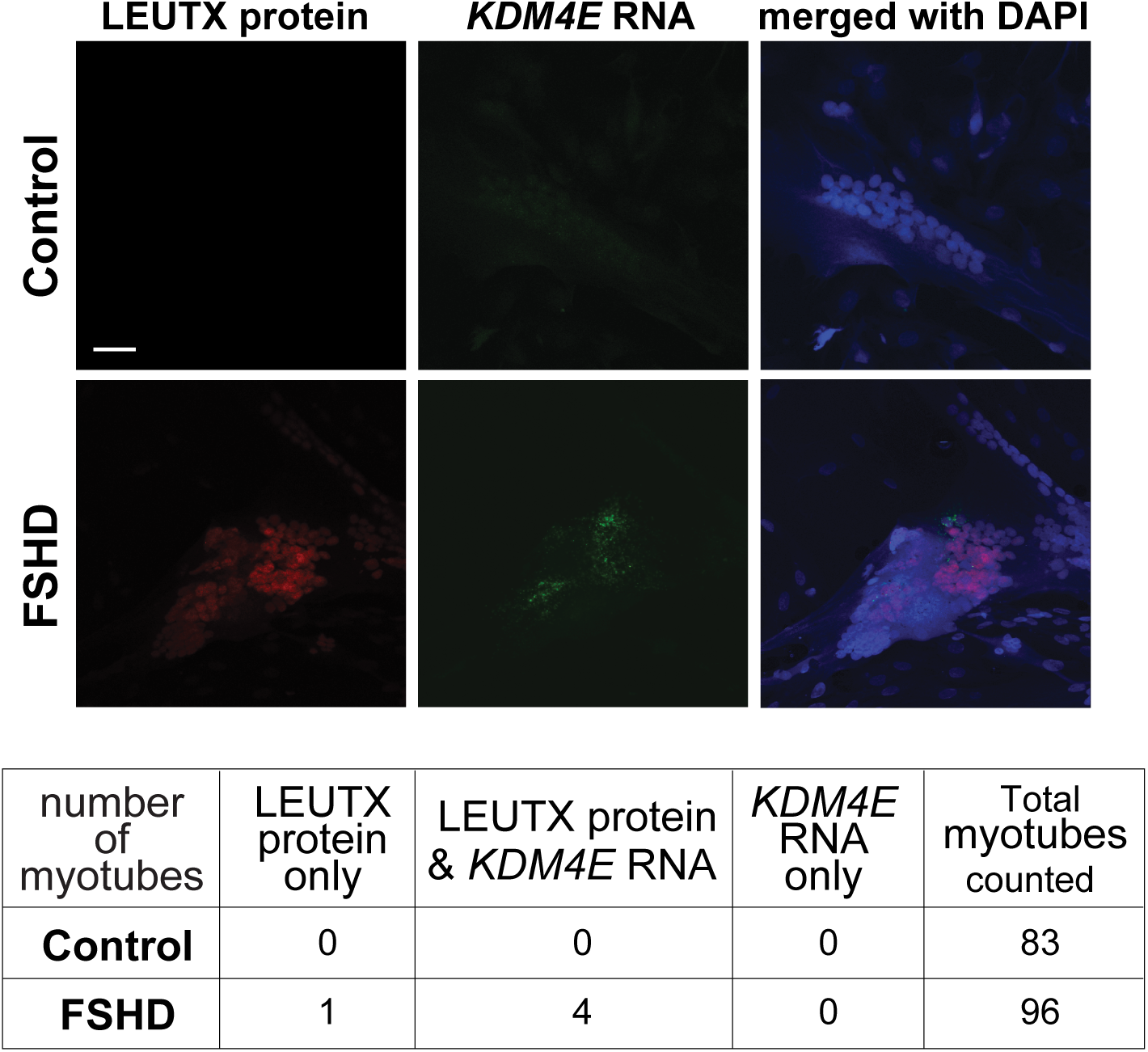
Immunofluorescence staining of LEUTX protein and *KDM4E* RNAScope on differentiation day 6 in FSHD immortalized myotubes. All observed *KDM4E* transcripts colocalized with LEUTX protein. Scale bar = 20 μm.

**Supplemental Figure S5.**
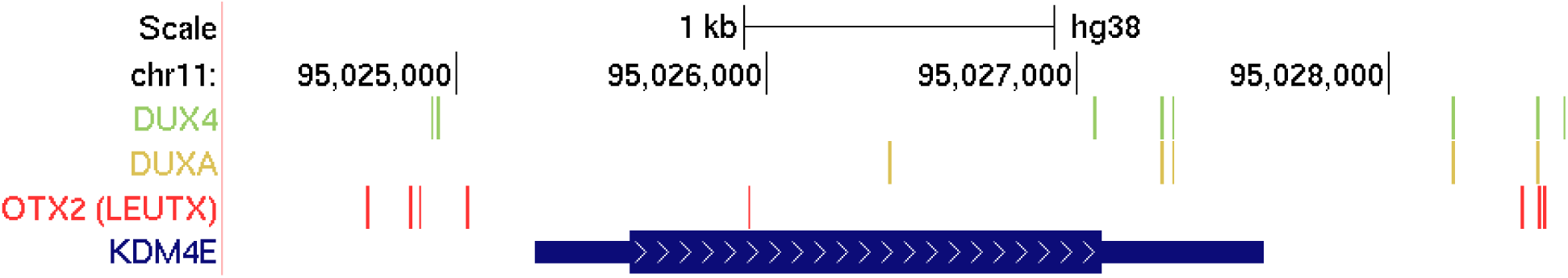
TF binding motifs at the promoter of *KDM4E*. Binding motifs for DUX4, DUXA and the putative LEUTX motif (OTX2) within 1 kb upstream and 0.5 kb downstream of the transcription start site (TSS) for *KDM4E*. We used binding motifs from HOCOMOCO v11 (1) for DUX4, DUXA and OTX2 as input into HOMER (version 4.10) using the scanMotifGenomeWide.pl command for hg38 (2). Since no binding motif was available for LEUTX, we used the motif for OTX2, another PRD-like homeobox TF with a similar homeodomain, which was experimentally validated to function as LEUTX binding site (3). Visualization was done on the UCSC genome browser (4) using GENCODE v28 for the *KDM4E* gene model.

